# Systematic Review of the Research on Morphological Modularity

**DOI:** 10.1101/027144

**Authors:** Borja Esteve-Altava

## Abstract

The modular organization of the phenotype is an emergent property that derives from a semi-independent relation of body parts in their inheritance, development, function, and evolution. Understanding the modular organization of living beings is essential to understand the evolvability and plasticity of organismal form, and how morphological variation is structured during evolution and development. For this reason, delimiting morphological modules and establishing the factors originating them is a lively field of inquiry in biology today. However, unifying the results of the available body of knowledge is a challenge because of the large number of species studied and the disparity of morphological system, from the flower of angiosperms to the wing of insects and the head of primates (to name a few). The specific factors behind each pattern of modularity and the methods used to identify vary widely as well. This review summarizes more than 190 original research articles since 1958, in order to get a quantitative appraisal on what is studied, how is studied, and how results are explained. The results reveal an heterogeneous picture, where some taxa, systems, and approaches are over-studied, while others receive minor attention; other major trends and gaps in the study of morphological modularity through time are also discussed. In sum, this systematic review seeks to offer an objective view of this research field and highlight future research niches.

## 1. INTRODUCTION

Modularity is a ubiquitous property of natural complex systems that emerges at all hierarchical levels of organization (Simon 1962; Callebaut 2005). The organization of the phenotype in modules is the result of the interplay between developmental processes, genetically and epigenetically controlled, and the functioning of morphological structures in its environment through time (Pigliucci and Preston 2004; Schlosser and Wagner 2004; Callebaut and Rasskin-Gutman 2005). A common developmental origin, an allometric relation of growth, a joint performance of a function, or a shared evolutionary history, are all examples of factors that can promote the integration of body parts into morphological modules. Moreover, modularity is a fundamental concept in biology that helps us to understand from the complexity of the genotype-phenotype map to the evolvability of the organismal form (von Dassow and Munro 1999; Bolker 2000; Müller 2007; Pavlicev and Hansen 2011). In the last years, many essays and narrative reviews have laid the foundation of an empirical research program on morphological modularity based on developmental, ecological, and evolutionary mechanisms (e.g. Raff and Raff 2000; Schlosser 2002; Pigliucci 2003; Wagner et al. 2007; Klingenberg 2008; Kuratani 2009; Murren 2012; Goswami et al. 2014; Klingenberg 2014; Rasskin-Gutman and Esteve-Altava 2014). A common take home message in all these studies is that a deeper insight of the modular organization of living beings is essential to understand the development and evolution of form. What we lack is a panoramic view of how we have approached the study of morphological modularity to date that helps us to identify the basis of our current knowledge and the gaps that will guide future research.

### 1.1. A Minimum Definition of Morphological Module and Integration

One of the problems of bringing together different type of morphological studies is that the definitions of module and of integration change depending on the level of organization studied and, most important, the type of factor guiding the study (see Eble 2005 for a conceptual review). Cheverud (1996) describes four types of factors of integration, that is, different types of interactions among body parts that integrate them within a same module: functional, developmental, genetic, and evolutionary interactions. The actual realization of these interactions might involve various processes and mechanisms, such as the physical distribution of forces, the diffusion of signaling molecules, or the inheritance of genetic regulatory networks controlling development. According to Cheverud, at an individual level, function integrates parts that perform the same or related tasks and need to coordinate during performance; while development integrates parts that interact during their formation, including those controlled by a same genetic network. At a population level, genes integrate parts that are inherited together (often due to pleiotropy: a single gene affecting multiple parts); and finally, evolution integrates parts that evolve in a coordinate manner because they are inherited or selected together. For a review on how genetic, developmental, functional, and evolutionary modules relate to each other see Klingenberg (2008). Other authors have proposed additional or complementary factors of integration. For example, Wagner and Altenberg (1996) introduced the operational concept of the variational module as correlated sets of traits that arise from variation in processes that affect those traits to a greater extent than others. In addition, Chernoff and Magwene (1999) argued that the organization of parts in the body integrates those parts that share structural relationships due to geometric and/or topological interactions. In his essay on the conceptual basis of morphological modularity, Eble (2005) introduced a distinction between variational modules *(sensu* Wagner and Altenberg), which are used to study how morphological parts co-vary during evolution and/or ontogeny, and organizational modules, which would capture the structural relations among body parts in individual organisms (e.g. Esteve-Altava et al. 2011). This distinction is essential in order to understand that, despite the mainstream view in morphometric studies, not all morphological modules need to relate to a covariance structure of shape; if that were the case, we would need additional studies to find the causes. For example, morphological modules in the human brain identified by its functional activity (e.g. using fMRI) are not necessarily linked to a pattern of covariation of shape (but see Gómez-Robles et al. 2014). More recently, Mitteroecker and Bookstein (2008) have distinguished between global factors that keep the cohesion of a morphological system (i.e. integrate the whole phenotype), and local factors that provide the internal cohesion of its modules (i.e. parcellate the whole phenotype). This distinction is important because, indeed, it is unknown whether each type of factor described above (e.g. function, development, genes) is uniquely related to the formation of morphological modules or the integration of the whole phenotype. In fact, the hierarchical nature of phenotypes suggests that the effect of a particular factor in the integration or parcellation of a body part depends on the scale this factor acts on (Bastir 2008).

In its simplest form, a morphological module is a group of body parts that are more integrated between them than to other parts outside the group (Eble 2005). Integration arises as a direct consequence of the number and strength of interactions, regardless of how we define interaction (Eble 2005). This minimum definition of morphological module uses the concepts *integration* and *interaction* vaguely on purpose: in order to be more inclusive. Because it makes no reference to why or how integration between parts emerges and varies (or the source of this integration), it applies to a wide range of morphological systems. In fact, by replacing “body parts” with “elements of a system” this definition applies even to non-biological systems (Simon, 1962). The concept of body part has also a broad sense to accommodate semi-independent structures (e.g. head, limbs) and anatomical units within a larger structure (e.g. bones of a skull, petals of a flower), as well as individual traits and morphometric measures, including their proxies (e.g. landmarks coordinates, distances). Notice that this minimum definition of morphological modularity is wider than traditional definition of morphological integration and modularity as related to a structure of covariation of shape and size (Terentjev 1931; Olson and Miller 1958), but it applies also to other sorts of morphological information (e.g. proportions, connections, articulations and orientations; see Rasskin-Gutman 2003; Rasskin-Gutman and Buscalioni 2001).

By specifying the factors behind integration (or parcellation) and the meaning of body part, each particular study turns this minimum definition of morphological module into an operational definition; this is an essential step in any quantitative study of modularity. However, the purpose of this review is to bring together as many research studies as possible, regardless of the morphological system, methodology, and factors of integration used. For this reason, the use of morphological modularity in this review will henceforth refer to the minimum definition of module, unless otherwise stated in each particular case.

### 1.2. On Morphological Modularity and Methodology

There are different methods available to validate morphological modules, from those derived from the quantification of variation in size and shape (e.g. multivariate analysis of linear distances or geometric morphometrics) to those based on the quantification of functional, developmental, and genetic interactions (Eble 2005). In the last 20 years, landmark-based geometric morphometrics has progressively replaced linear distances to measure morphological integration and modularity among body parts (Adams et al. 2013). One reason for this methodological shift is the advent of more sophisticated software for image digitization and statistical analysis. In addition, new developments in network theory have been applied to the study of the complex structural and functional integration of the human brain (Sporns 2011). More recently, methodological approaches that seemed to be restricted to the study of some type of morphological structures are starting to be transferred to the study of other structures. For example, geometric morphometrics has been used recently to analyze modularity in the brain (Gómez-Robles et al. 2014), while network analysis has been applied to find the morphological modules of the head (Esteve-Altava et al. 2013; Esteve-Altava et al. 2015).

### 1.3. Open Questions

This review compiles and evaluates 191 original research articles that explicitly studied the presence and/or validation of morphological modules in animals and plants. Through a systematic quantification of the materials of study, the methodological approaches, and results obtained in these research articles, this review seeks to answer the following specific questions:

1. What are the sources of our knowledge on morphological modularity and where are the gaps (if any) ?

a. Is morphological modularity ubiquitous in all living beings?
b. Is morphological modularity equally pervasive in all body parts?
c. Are there biases in the study of morphological modularity?
2. How do we study morphological modularity?

a. What biological criteria do we use to propose hypotheses of modularity?
b. What methods do we use to test these hypotheses?
c. Do the same biological factors explain modularity and integration patterns?
3. And finally, what are the most representative examples of morphological modules in different organisms and morphological systems?

## 2. METHODS

### 2.1. Gathering of Original Research Studies

The studies reviewed comprise only peer-reviewed original research articles that explicitly assess morphological modularity. I searched in Google Scholar for articles that include in their title at least one of the following keywords: functional integration, genetic integration, modular evolution, modularity, morphological integration, mosaic evolution, ontogenetic integration, phenotypic integration, or pleiades. The year of publication of the landmark book *Morphological Integration* by Olson and Miller in 1958 was used to set the beginning of the search. To not exceed the limit of 1,000 entries in Google Scholar, I performed a separate search for each individual year between 1958 and June 2015.

Google Scholar has a proven high coverage--100% for medical systematic reviews--despite its low precision (i.e. many entries retrieved are irrelevant) (Gehanno et al. 2013). This means that Google Scholar can be used alone in systematic reviews without missing any relevant reference. Unfortunately, this searching method does not avoid the fact that some relevant research articles might be excluded from the sample if their titles do not include the above keywords. The aim of the search would be then to retrieve a sufficient amount studies that represent the general practice of the research in morphological modularity. That said, the dataset that evaluates the research articles has been designed to include new items as they are available. The statistical exploration performed on the dataset is also completely reproducible (and the code openly available) in order to allow a future analysis with new items. The whole project is accessible at https://osf.io/h4ka7/.

The search retrieved more than 5,500 results of which 610 seemed to match the selection criteria. I updated and searched the full text of each potential entry using EndNote X7 through several institutional journal subscriptions (Universitat Jaume I, Universitat de València, Universitat Autònoma de Barcelona, Royal Veterinary College, and Howard University), public repositories, personal webpages, and personal requests. A total of 191 articles were selected for review after excluding duplicate entries, non-original-research articles (e.g. books, chapters, essay, and other reviews), and articles off topic that did not assessed morphological modularity. Exceptionally, two articles were included twice in the dataset (Magwene 2001; Klingenberg 2009), because they comprised two independent studies, each one requiring a separate evaluation.

### 2.2. Evaluation of Original Research Studies

For each entry in the dataset I collected the following information: authors, year of publication, title, field of the journal of publication, taxa (genus, family, order, class, phylum, and kingdom), system of study, type of material (fossil and/or living), type of tissue (hard and/or soft), scale of the study (macroevolutionary, microevolutionary, ontogenetic, or any combination of these), number of null hypotheses tested, specific hypotheses tested, criteria used to define these hypotheses (anatomy, development, function, genetics, growth, origin, shape, size, timing, and/or other criteria), methods used to identify or validate morphological modules (biomechanical performance, geometric morphometrics, heterochrony analysis, linear metrics, network analysis, qualitative description, quantitative trait loci, and/or other methods), results obtained (number of studies validating an hypothesis, number of studies reporting integration, and specific results of modularity), and factors argued for these results (developmental, environmental, functional, genetic, growth, phylogeny, topological, and/or other factors).

### 2.3. Details of the Coding of Variables

- *Authors:* the names of the authors were recorded to identify the study.
- *Year of publication*: the year of publication was coded using four digits (e g. 2015).
- *Title*: the title of the publication was recorded to identify the study.
- *Field of the journal of publication*: the field was assigned according to journal description using the following categories: anatomy, anthropology, general biology, botany, cell biology, development, ecology, evo-devo, evolutionary ecology, evolution, generalist, genetics, medicine, neurosciences, paleobiology, physics, physiology, and zoology.
- *Taxa:* the genus, family, order, class, phylum, and kingdom of the specimens used in the study were coded as different variables. For simplicity, only the lowest rank that includes all the specimens of the study was coded. For example, a study comparing primates and rodents was coded as class Mammalia, while order, family, and genus were coded as NA.
- *System of study*: a descriptive label of the system used in the study, for example, skull, body parts (i.e. comparison among different structures such as limbs and heads), flower, or brain. More inclusive labels were used when various components are studied; for example, a study analyzing the skull together with the brain or with attached muscles would be coded as head.
- *Type of material:* this was coded as being fossil or living, while the label both was used when the sample included extinct and extant species.
- *Type of tissue:* only applied in vertebrates, hard tissue was used for studies analyzing skeletal and cartilaginous samples; soft tissue was used for tissues other than bones and cartilage (e.g. brain, muscles, organs); and both was used when the study included hard and soft tissues.
- *Scale of the study:* this variable codes for the temporal scale of the analysis. Micro-evolutionary refers to studies at the population level, including samples within only one species; macroevolutionary refers to studies comparing different species or higher taxa; ontogenetic refers to studies comparing different developmental stages; case study refers to studies analyzing only one specimen or a very small sample (n<5). Studies combining these scales were labeled accordingly (e.g. Micro/Macro, Onto/Macro).
- *Number of null hypotheses tested:* an integer, the number of hypotheses explicitly tested for the authors of the study.
- *Specific hypotheses tested* (if any): a description of the modules tested. The term integration is used when the hypothesis explicitly tested is the absence of modules.
- *Criteria used to define these hypotheses:* a list of different variables was evaluated and coded independently, anatomy, development, function, genetics, growth, origin, shape, size, timing, and others. The list captures the most frequent criteria identified in the dataset. For each variable, 1 indicates that authors used this criterion explicitly or implicitly as a source to make their hypotheses. Each variable is coded separately as 1 if more than one hypothesis is tested or one hypothesis is based on several of the criteria listed. For example, anatomy and function would be coded as 1 in a study that test a hypothesis of modularity based on the anatomical structure of the system and its function.
- *Methods used to identify or validate morphological modules*: a list of different variables was evaluated independently, biomechanical performance, geometric morphometrics, heterochrony analysis, linear metrics, network analysis, qualitative description, quantitative trait loci, and others. The list captures the most frequent criteria identified in the dataset. Each variable is coded separately as 1 if authors used this family of methods. For example, a study that uses geometrics morphometrics and quantitative trait loci would be coded as 1 in both variables.
- *Results obtained:* three variables capturing separately, whether the study validates a proposed hypothesis (1) or not (0), whether the study reports integration (1) (i.e. absence of modularity) or not (0), and the specific result of the study regarding the presence modules (if any).
- *Factors argued for these results:* a list of different variables was evaluated independently, developmental, environmental, functional, genetic, growth, phylogeny, topological, and others. The list captures the most frequent factors identified in the dataset. Each variable is coded separately as 1 if authors used this type of factor to support their results. For example, a study that reports that development is the sole cause of a modular organization has 1 in the variable development. Multiple factors are possible.

## 3. RESULTS AND DISCUSSION

The empirical study of morphological modularity has grown exponentially in the last 25 years. Almost half of these studies were published in journals with a strong focus on evolutionary biology, while the other half were published either in specialist or in generalist journals (Fig. 1). This publishing pattern highlights a rising interest on morphological modularity in the biological community as a whole, and in particular, of the impact of modularity to understand the development and evolution of living beings. The following sections summarize and discuss the main results of the systematic quantification of the reviewed research articles.

**Figure 1.**
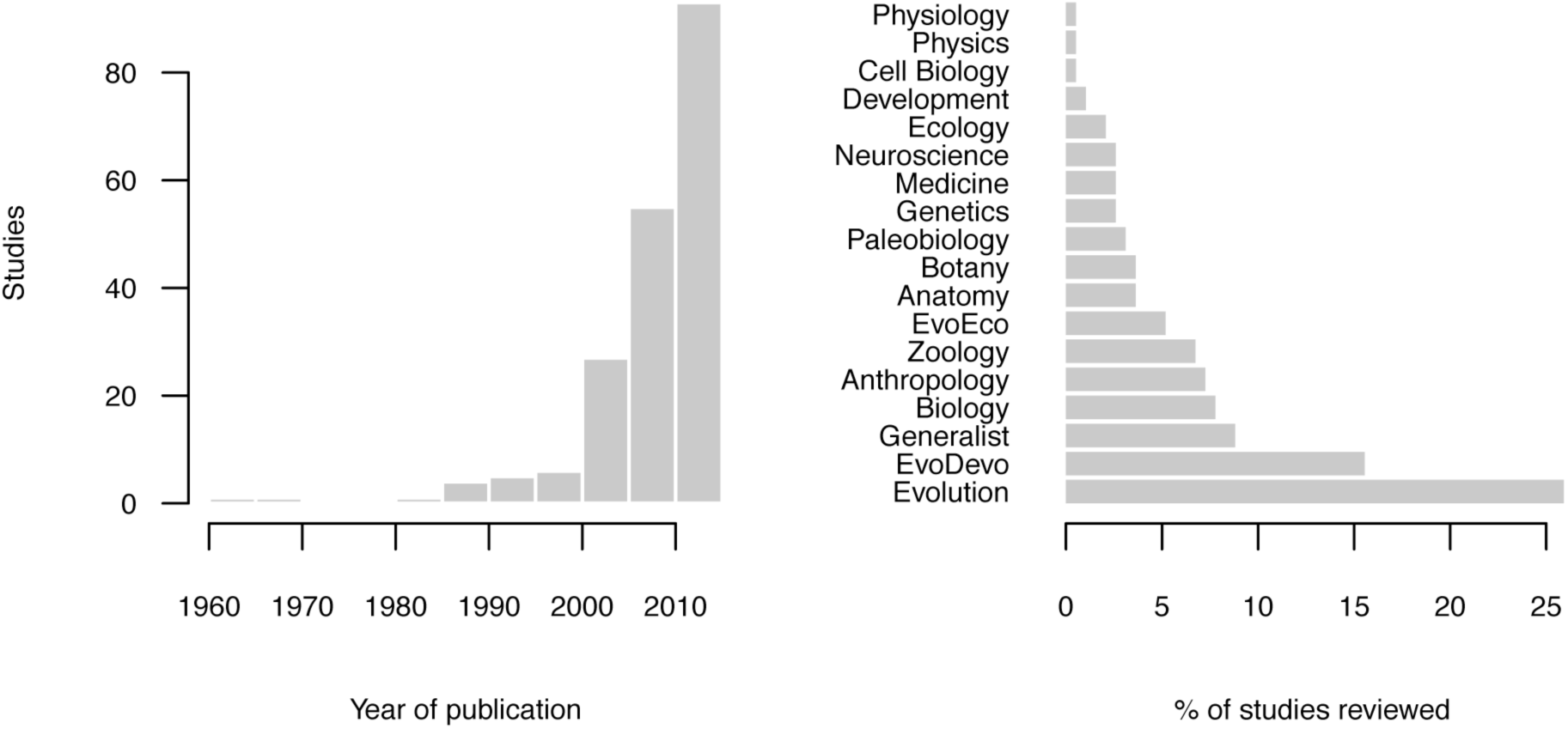
Number of publications about morphological modularity from 1959 to 2015 and the top-10 research fields of publication of the literature reviewed. Notice that for 2015 the sample only includes articles published in the first six months.

### 3.1. Our Knowledge of Morphological Modularity (and its Gaps)

Most of our knowledge about morphological modularity comes from the study of mammals (61% of the studies), with a strong focus on primates and rodents (Fig. 2, *top*). At the genus level, *Homo* (20%) and *Mus* (12%) are the predominant taxa. Presumably, the preference for these groups is due to a particular interest in the morphological evolution of our own species, as well as the use of mice as a model species. More surprising is the low number of studies that focus on the modularity of plants (11%), despite there is a long tradition in the study of phenotypic integration in this group (e.g. Berg 1960; Pigliucci et al. 1991; Diggle 2014). The arthropods are also underrepresented (10%), despite having a segmented body plan (composed of tagmata or metameres), which makes this group of animals particularly interesting for inquiring about modularity at a morphological level (e.g. Yang 2001; Yang and Abouheif 2011; Molet et al. 2012). Unfortunately, research is marginal in other phyla that are often referred as being modular organisms, such as sponges and fungus, and inexistent in some phyla. This bias in the taxa studied raises some doubts about the ubiquity of modularity in all multicellular, eukaryotic organisms. Although the presence of a modular organization in all organisms has a solid conceptual and empirical foundation, our understanding would benefit from including these underrepresented taxa in future studies on morphological modularity.

**Figure 2.**
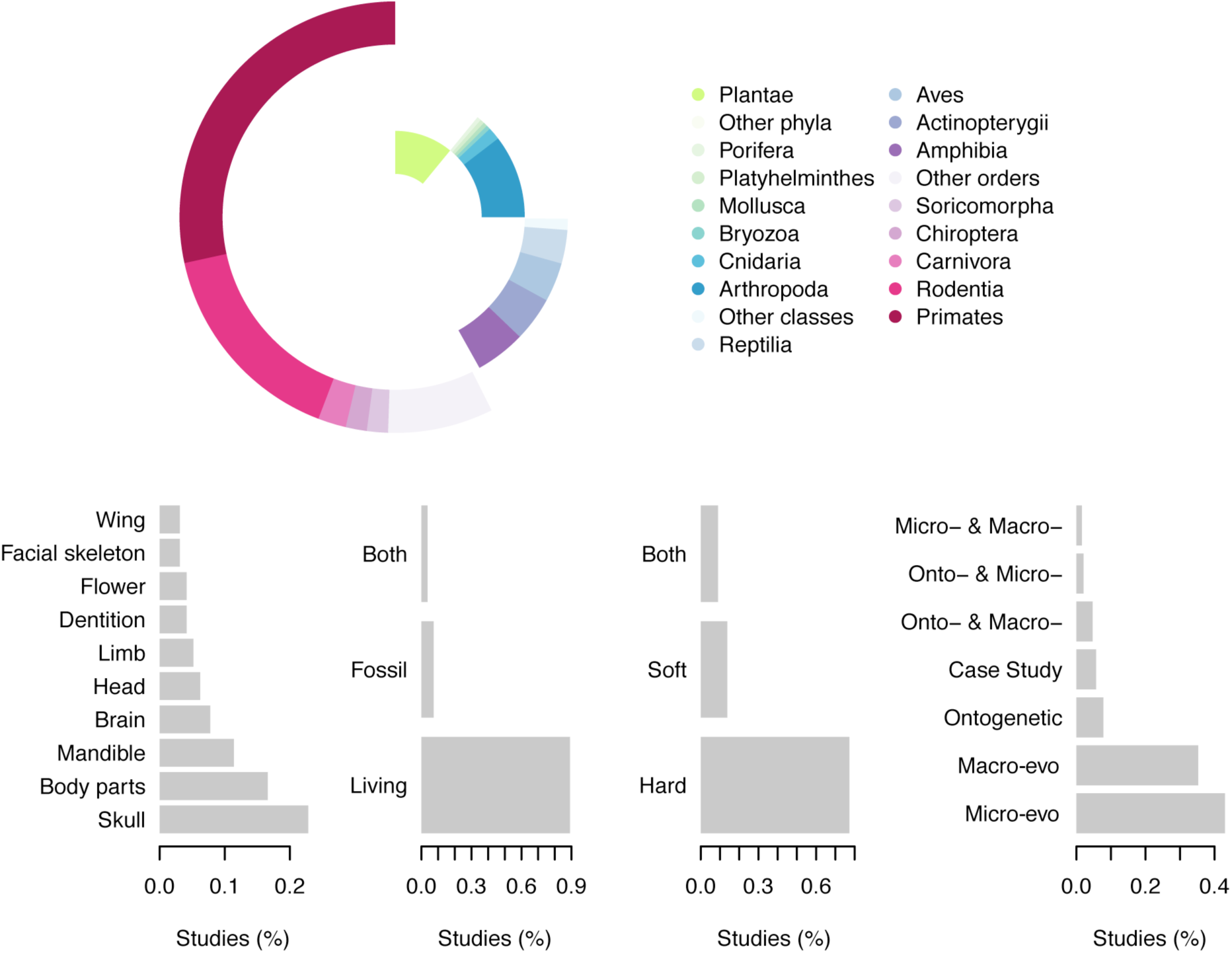
Patterns in the objects of studies of morphological modularity. *Top,* schema showing the proportion of taxa studied in the literature reviewed, with emphasis on most commonly studied taxa. *Bottom,* patterns in the design of studies in the literature reviewed: System, tissue, material, and scale.

The head of vertebrates is the most studied morphological structure (Fig. 2, *bottom*). However, the majority of these studies focus on separate components of the head, such as the skull (23%), the mandible (11%), the brain (8%), or the dentition (4%); while the head as a whole is considered only in a few cases (e.g. Hünemeier et al. 2014; Tsuboi et al. 2014). The 17% of the studies compared morphological integration within and among different parts of the body (i.e. considering them as individual modules), such as cranial vs. postcranial skeleton, forelimb vs. hindlimb, and flowers vs. leaves. Finally, other morphological structures that are commonly acknowledged for having well-delimited modules are relatively underrepresented, such as the limb of tetrapods (5%), the flower of angiosperms (4%), and the wing of insects (3%) (e.g. Klingenberg and Zaklan 2000; Hamrick 2012; Diggle 2014), have attracted relatively less attention than the head. Delimiting morphological modules is probably trickier (and more challenging) in the head than in other parts of the body, because of the many overlapping developmental and functional interactions among the structures of the head (Lieberman 2011a), which may obscure the patterns of covariation among parts (Hallgrimsson et al. 2009). As a consequence, the head is probably the morphological system for which more different modular hypotheses have been proposed.

The literature reviewed shows noteworthy biases toward overrepresentation of hard tissues, extant species, and evolutionary scale (Fig. 2, *bottom).* Notably, only 14% of the studies in vertebrates analyze soft tissues, including the brain. This bias contrasts, for example, with the importance placed on developmental and functional relations between hard and soft tissues to explain the morphological evolution of vertebrates (Diogo and Wood 2013; Richtsmeier and Flaherty 2013). Hard tissues are sometimes regarded as more useful for practical reasons (e.g. easier to work with, fossil comparisons) or more evolutionary stable (i.e. showing less homoplasy). Exhaustive studies on the evolution of the muscular system in primates suggest that the latter argument lacks of empirical support (e.g. Diogo and Wood 2011, 2013). The prevalence of hard tissues due to their use in fossil studies or comparisons does not hold neither, because only 11% of studies use fossil materials (alone, 7%, or in combination with extant species, 4%). Finally, most of the studies reviewed follow micro- or macroevolutionary evolutionary scales (80%), while only 15% of the studies follow exclusively or partially an ontogenetic scale, despite the importance of development as a primary source of morphological modularity. These results suggest that we know relatively little about morphological modularity of systems made only of soft tissues, or of systems combining hard and soft tissues. The study of modularity in soft and hard/soft structures such as the brain, brain/skull, and bones/muscles is growing in recent times (e.g. Richtsmeier et al. 2006; Gómez-Robles et al. 2014; Esteve-Altava et al. 2015). This is an endeavor worth enduring according to the results reported here.

### 3.2. Trends in the Study of Morphological Modularity

A typical research study of morphological modularity follows a two-step approach: first, proposing an hypothesis of modules according to functional, developmental, genetic, or evolutionary criteria; and then, testing this hypothesis using quantitative methods. For example, in studies using morphometric methods (the most common; see below) a hypothesis of modularity is validated if it conforms to an observed pattern of variational modularity (Klingenberg 2008). Accordingly, we refer to functional, developmental, genetic, and evolutionary modules depending on the integrating factors at play. Studies that do not test any hypothesis take an exploratory approach, for example, using different methods to look for morphological modules with no previous assumptions (e.g. Magwene 2001; Esteve-Altava et al. 2013).

Function and development are the two most common criteria used to define morphological modules, but their predominance vary in animals and plants (Fig. 3, *top*). In animals, development and functional criteria are used alike (33%) to generate hypothesis of morphological modules. In plants, function (59%) is more predominant than development (23%) to delimit modules. This result suggest that studies in plants are more interested in the functional relation among body parts, which has important ecological consequences. However, an historical reason can be argued as well to explain this difference. Animal studies often refer to Olson and Miller’s book *Morphological Integration* (Olson and Miller 1958), as well as Cheverud’s works published during the 1980s and 1990s analyzing patterns of developmental and functional integration in the cranial skeleton (Cheverud 1996). In contrast, plant studies usually refer to the work of Berg on *The Ecological Significance of Correlation Pleiades* (Berg 1960), where she proposed a modular division of angiosperms based on reproductive and vegetative functional criteria. Different traditions (or inertia) in the use of developmental and functional/ecological factors in determining the form of animals and plants might explain the difference observed also in the criteria used to delimit morphological modules.

**Figure 3.**
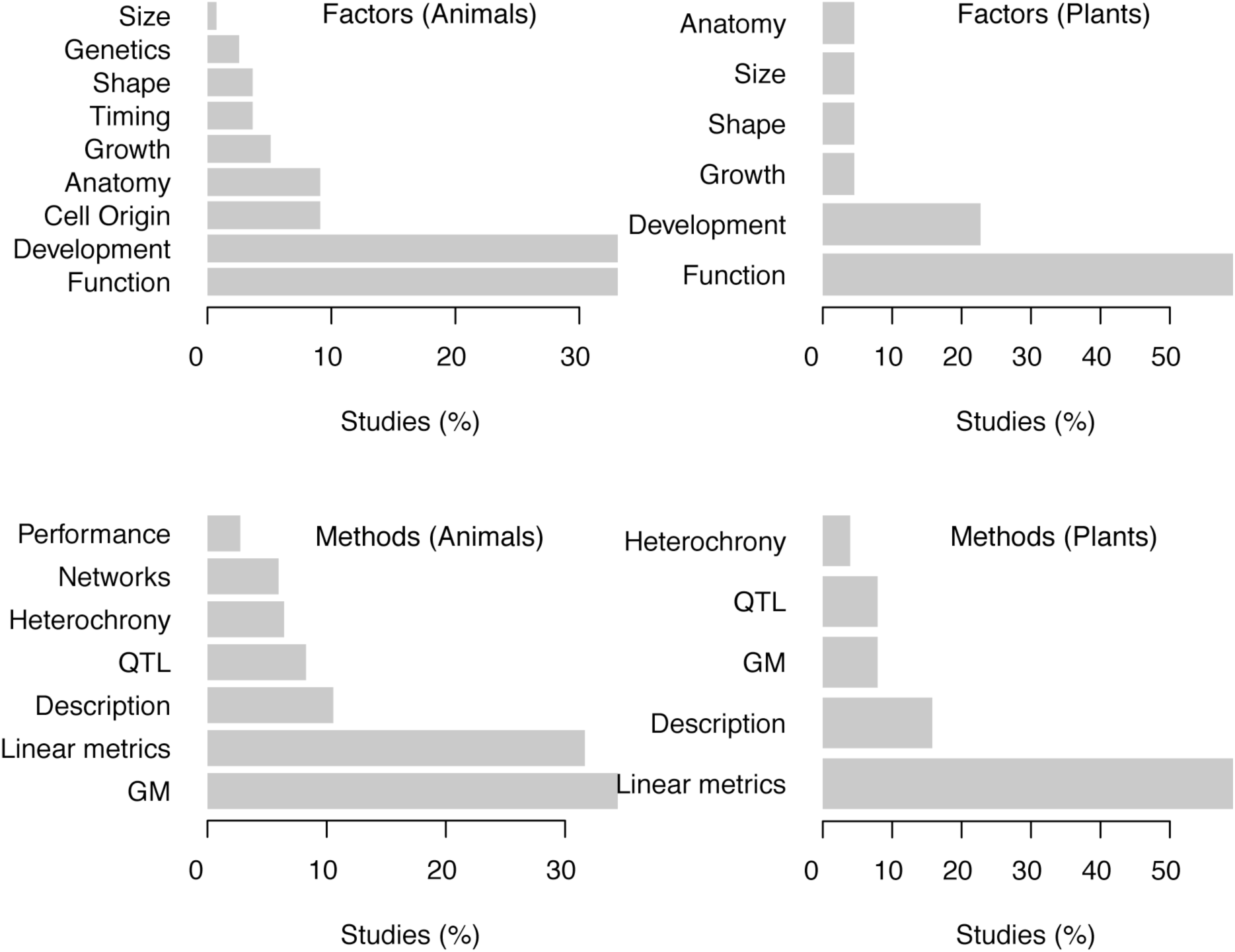
Patterns in the design of studies of morphological modularity. *Top,* criteria used to delimit morphological modules in animals and plants. *Bottom,* methodological approach.

The most common methods used in the studies reviewed are related to the quantification of size and shape, whereas description, quantitative trait loci, heterochrony, network analysis, and biomechanical performance are less common (Fig. 3, *bottom).* In animal studies, both geometric morphometrics (34%) and linear metrics (32%) methods are similarly frequent. In plant studies, linear metrics (62%) are much more frequent than geometric morphometrics (8%). This difference might be only a consequence of the more recent introduction of geometric morphometrics in plants than in animals. In the context of morphological modularity, the earliest studies using geometric morphometrics in animals date from the late 1990s (Monteiro and Abe 1997; Adams 1998), while in plants date from the early 2010s (Klingenberg et al. 2012). The novel use of network theory in morphology is particularly interesting. Network analysis is common in neurosciences to identify modules, for example, using community detection algorithms (Fortunato 2010), because the very structure of the brain, as a web of neurons, is readily modeled as a network (Sporns 2011). However, network methods have been also applied more recently to study the morphological modularity of the human head: the nodes of the network represent the bones and muscles of the head, connected through their physical interactions (e.g. Esteve-Altava et al. 2013; Esteve-Altava and Rasskin-Gutman 2015; Esteve-Altava et al. 2015). This novel use of network models in anatomy should not be confused with the use of *“graph modeling”,* a method that uses graphs to represent phenotypic correlation of morphometric traits or landmarks in statistical analyses (e.g. Magwene 2001, 2008; Zelditch et al. 2009). Notably, graph modeling of morphometric traits correlation has been recently used together with network analysis to study the modules of the wing of some insects (Suzuki 2013).

Independently of the method used, only 39% of the studies reviewed validated at least one of the hypothesis proposed; the remaining 61% either rejected all hypotheses or found an unexpected result, not considered in any initial hypothesis. In total, 73% of the studies reported a pattern of modularity in the system studied, while the other 27% reported that whole-system integration is stronger than modularity (i.e. modules were not identifiable or delimited). The four leading factors to explain these results were functional, developmental, genetic, and environmental. Their relative importance varies however between animals and plants, as well as between studies reporting a results of modularity and of integration (Table 1). In animals, function and development are the two most frequent factors in explaining both modularity and integration, followed by genetic and environmental factors. Thus, there is not an association between the type of factor and its effect on the formation of modules or the integration of the whole system. In contrast, in plants external factors (functional and environmental) are dominant in explaining modularity, while internal factors (genetic and developmental) are slightly more dominant in explaining integration.

**Table 1.**
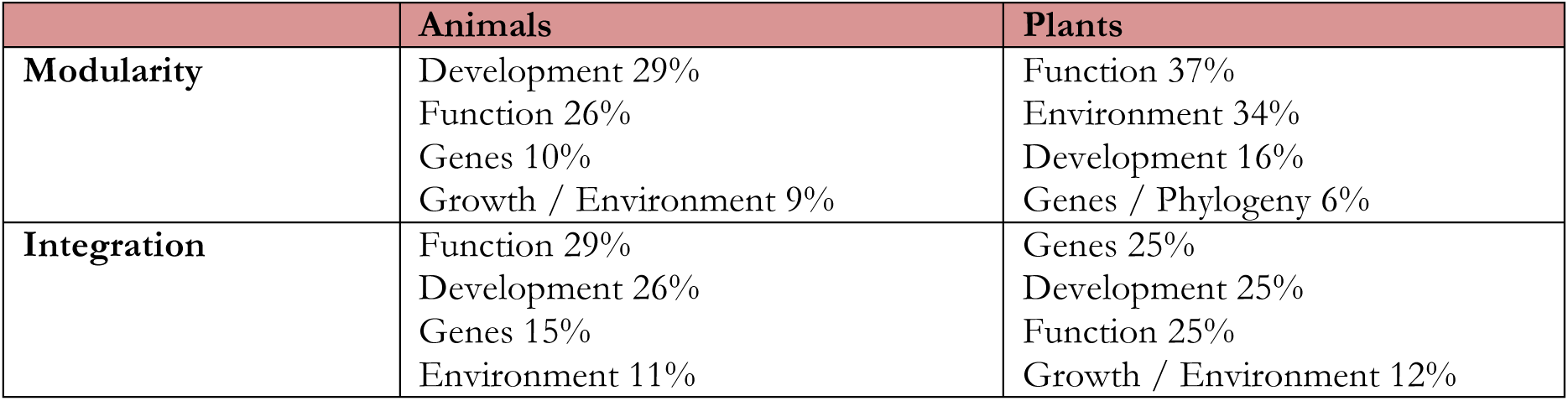
Factors most frequently used to explain results of studies of morphological modularity.

|  | **Animals** | **Plants** |
| --- | --- | --- |
| **Modularity** | Development 29%<br>Function 26%<br>Genes 10%<br>Growth / Environment 9% | Function 37%<br>Environment 34%<br>Development 16%<br>Genes / Phylogeny 6% |
| **Integration** | Function 29%<br>Development 26%<br>Genes 15%<br>Environment 11% | Genes 25%<br>Development 25%<br>Function 25%<br>Growth / Environment 12% |

### 3.3. Examples of Morphological Modules

This section summarizes the morphological modules reported in the literature reviewed for some of the most popular systems: the vertebrate skeleton (skull, mandible and limbs), the primate brain, the insect wing, and the body of angiosperms. Note that this is not an exhaustive list or an evaluation of the merits of each hypothesis of morphological modularity proposed; rather this is a glimpse of the most common modules proposed for each system to date. For this reason, each subsection includes references to more specific literature reviews.

#### The skull

The mammalian skull is the most popular structure studied in the literature reviewed (Fig. 4A). The skull is generally divided into various modules, hierarchically nested by developmental and functional interactions (e.g. Makedonska 2014). Often the skull is divided into orofacial, cranial base, and cranial vault modules, which is an instance of the classic division into face and neurocranium (Moore 1981). However, the precise boundaries among these modules vary in each study, depending on the species analyzed, the materials available, and the methodological approach. For example, studies using geometric morphometrics usually delimit the face-neurocranium boundary placing landmarks within the frontal, sphenoid, and zygomatic bones (e.g. Bastir and Rosas 2005; Goswami 2006; Mitteroecker and Bookstein 2008). In contrast, studies using anatomical network analysis placed the frontal within the facial module, whereas the sphenoid and the zygomatics are placed within the neurocranium (Esteve-Altava et al. 2013; Esteve-Altava and Rasskin-Gutman 2015). An also common modularity hypothesis divides the skull into oral, nasal, orbital, zygomatic, base, vault modules (Cheverud 1982), or even smaller modules (e.g. Makedonska 2014). This modularity hypothesis derives from Moss’ *functional matrix hypothesis* (Moss and Young 1960), which states that skeletal units (i.e. skull modules) develop and evolve in response to the functional demands of surrounding soft tissues and cavities. Although the existence of these functional matrices is generally accepted, some studies have also moderated their relative importance in shaping skull modularity and morphological variation (e.g. Lieberman 2011a, 2011b; Esteve-Altava and Rasskin-Gutman 2014). However, we lack enough studies that evaluate the modularity of the skull in relation to, and together with, its surrounding soft tissues (but see Richtsmeier et al. 2006; Esteve-Altava et al. 2015 for examples toward this direction).

**Figure 4.**
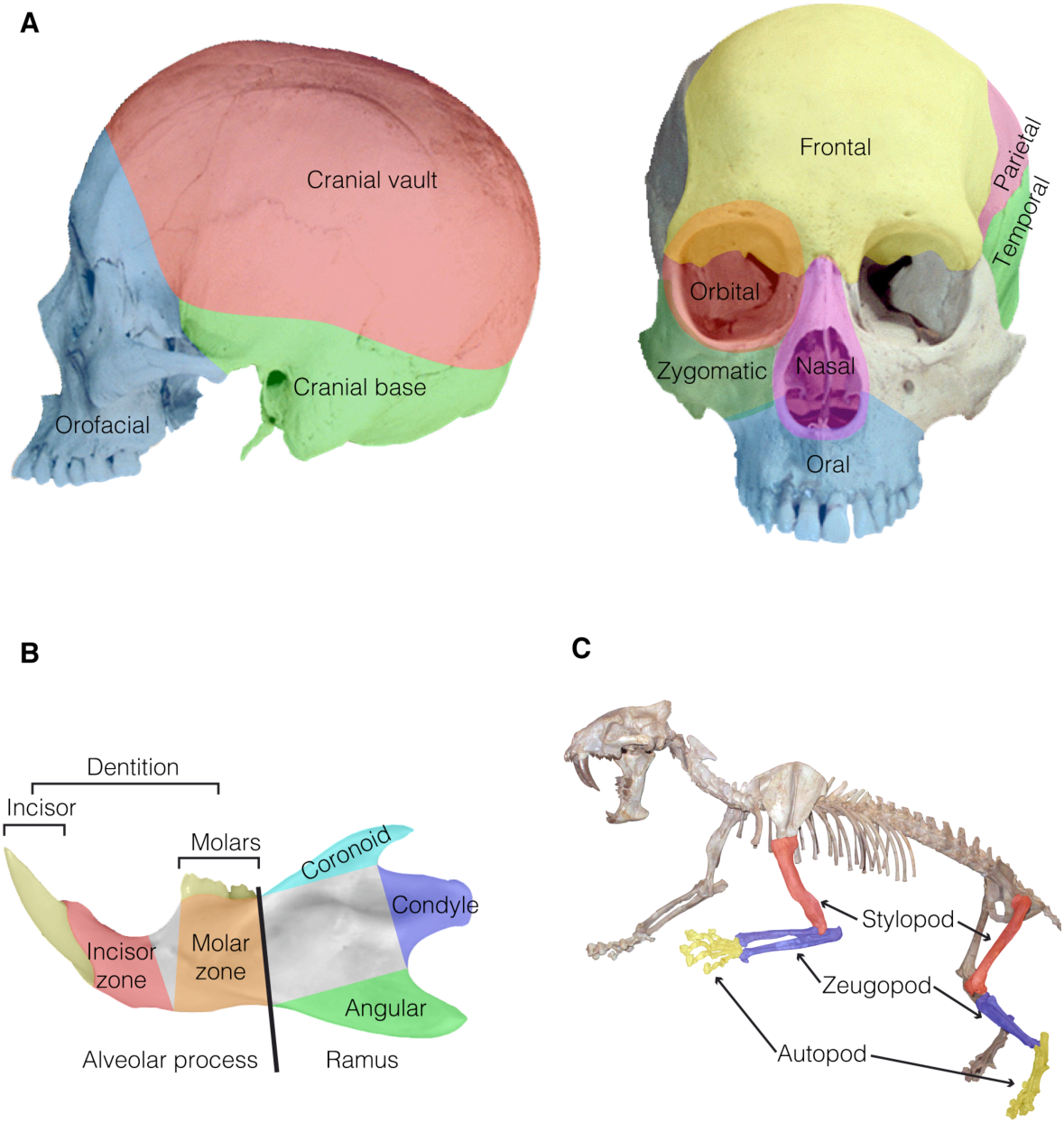
Morphological modules proposed for the mammalian skeleton. (A) *Left,* classical division of the skull in three modules: face, cranial base, and vault. *Right,* modules proposed by Cheverud (1982) according to the Functional Matrix Hypothesis (Moss and Young 1960): each module comprises those skeletal units affected by the formation and functioning of each functional matrix of the head (e.g. nasal, oral and brain). (B) Modules of the mandible illustrated in *Apodemus sp.* (Rodentia: Mammalia). The alveolar process and the ramus are the two main divisions of the jawbone. Additionally, various regions of the mandible are considered morphological modules according to their development, forming from different ossification centers, as well as their function related with the attachment of muscles, the articulation, and the bearing of teeth. The dentition is often studied as a separated structure, with each teeth series as an independent module. (C) Modules of the tetrapod limb illustrated in *Hoplophoneus dakotensis* (Carnivora: Mammalia). The autopod, the zeugopod, and the stylopod are the three modules of the limb, with the girdle sometimes as an additional module. Image credits: original photograph of the human skull from eSkeletons.org; original photograph of the Hoplophoneus skeleton by Rama (Source: Wikimedia commons).

#### The mandible

Closely related functionally, developmentally, and topologically to the skull, the mandible is often studied as a separate morphological structure in studies on modularity, because it has its own pattern of modularity independently of the rest of the skull (Fig. 4B). According to developmental and genetic criteria, the mandible comprises two modules: an anterior alveolar region and an posterior condylar ramus (Klingenberg et al. 2003). This division is also functional because the alveolar module bears the dentition, while the condylar ramus articulates with the skull and serves as the insertion surface for many masticatory muscles. However, according to functional and evolutionary criteria the mandible can be further sub-divided into five modules: in the alveolar region, the tooth-bearing incisor and molar zones; and in the ramus region, the coronoid, condylar, and angular processes (e.g. Ehrich et al. 2003; Renaud et al. 2012). Moreover, the dentition can be considered as a module within the jawbone, or as a semi-autonomous system that is further divided into tooth-row modules and tooth modules (e.g. Labonne et al. 2014).

#### The postcranial skeleton

The study of modularity in the postcranial skeleton has focused mainly in the study of limbs (Fig. 4C). The limb is commonly divided according to functional and developmental criteria into the stylopod (humerus; femur), the zeugopod (ulna and radius; tibia and fibula), and the autopod (wrist and fingers; ankle and toes). Some studies approaches the modularity of limbs by comparing the patterns of morphological integration among these three units in a same limb, between left and right limbs, and between forelimb and hindlimb (e.g. Martín-Serra et al. 2015; see also Goswami et al. 2014). Less common is the study of the morphological modularity in a single bone, such as the scapula (Young 2004), the humerus (Arias-Martorell et al. 2014), and the tibia (Tallman et al. 2013).

#### The brain of primates

The morphological division of the brain is commonly related to the embryonic origin of each of its parts, such as the forebrain (telencephalon and diencephalon), the midbrain (mesencephalon), and the hindbrain (metencephalon and myelencephalon) (Redies and Puelles 2001). An alternative hierarchical division of the brain is also sometimes considered (Marrelec et al. 2008), in which the brain divides into left and right hemispheres, and these, in turn, into regions or lobes (frontal, parietal, temporal, and occipital). These units are also studied in relation to their functioning, as regions that interact in performing a given task (i.e. functional integration), in different normal and pathological conditions. In a broader context, Gómez-Robles and co-workers (2014) offer a comprehensive studies of the brain morphological modularity, by quantifying and comparing the morphological integration using geometric morphometrics of different divisions of the brain, according to functional, structural, and evolutionary and developmental criteria at larger and finer scales (Fig. 5).

**Figure 5.**
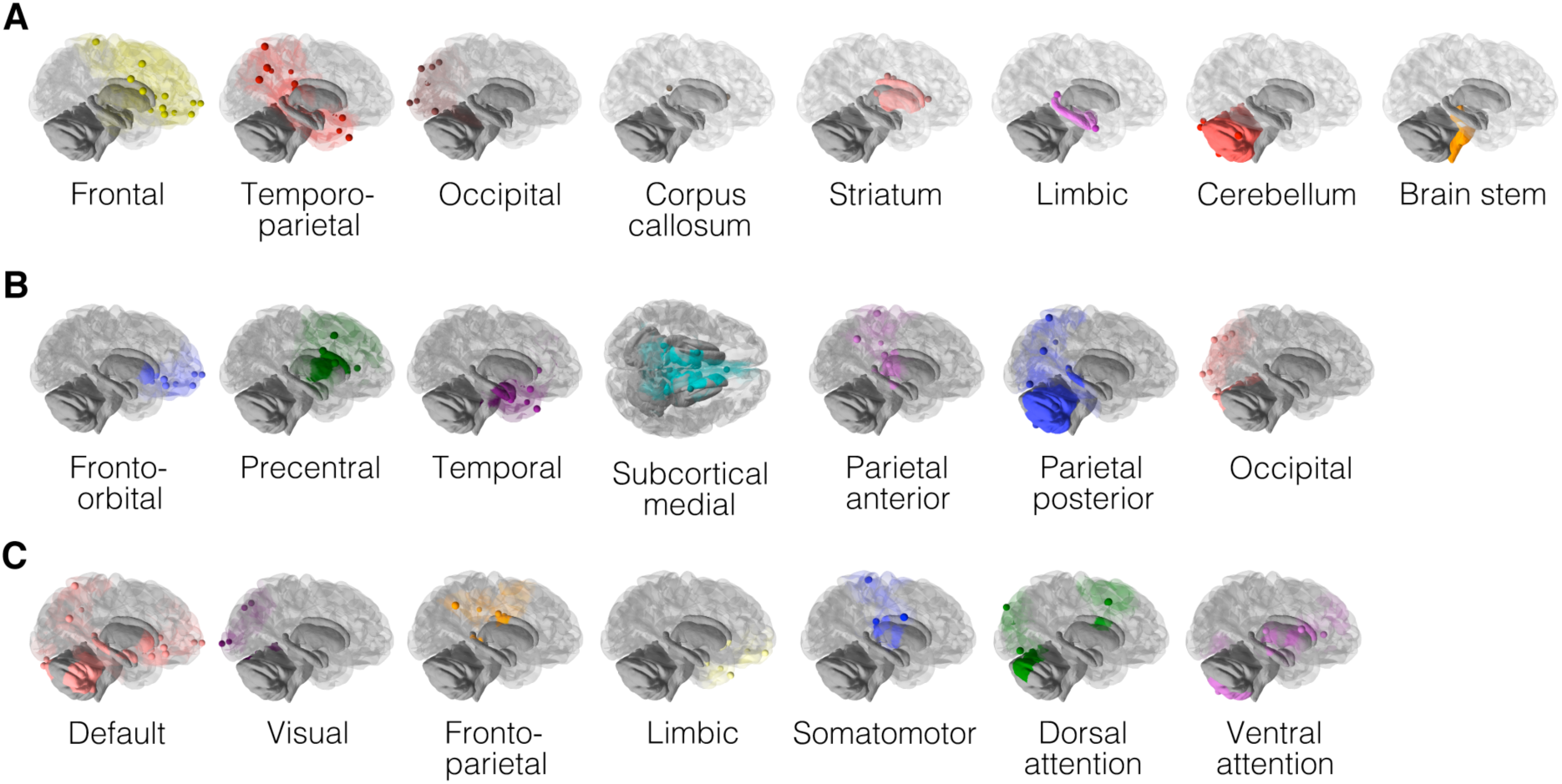
The modules of the brain illustrated in *Homo sapiens.* The image shows different modularity hypotheses of the brain according to (A) evolutionary and developmental criteria, (B) structural criteria, and (C) functional criteria. Modified with permission from Gómez-Robles et al. (2014) © NPG. (**Author Note**: permission to reproduce this figure is free of charge, the formal permission is needed only after the manuscript is accepted for publication.)

#### The wing of insects

Arthropods have a clearly recognizable modular body plan (Wagner 1990), and yet the studies reviewed of this group focus mainly on the wing of insects. The most common division in the insect’s wing is into two modules: one anterior and one posterior (Klingenberg et al. 2001) (Fig. 6A). In insects with two pairs of wings, the forewing and the hindwing have been reported also as being two different modules. Additionally, other geometric divisions, based on the wing patterns, have been also used to describe the wing of some insects (e.g. the nymphalid ground plan, see Suzuki 2013).

**Figure 6.**
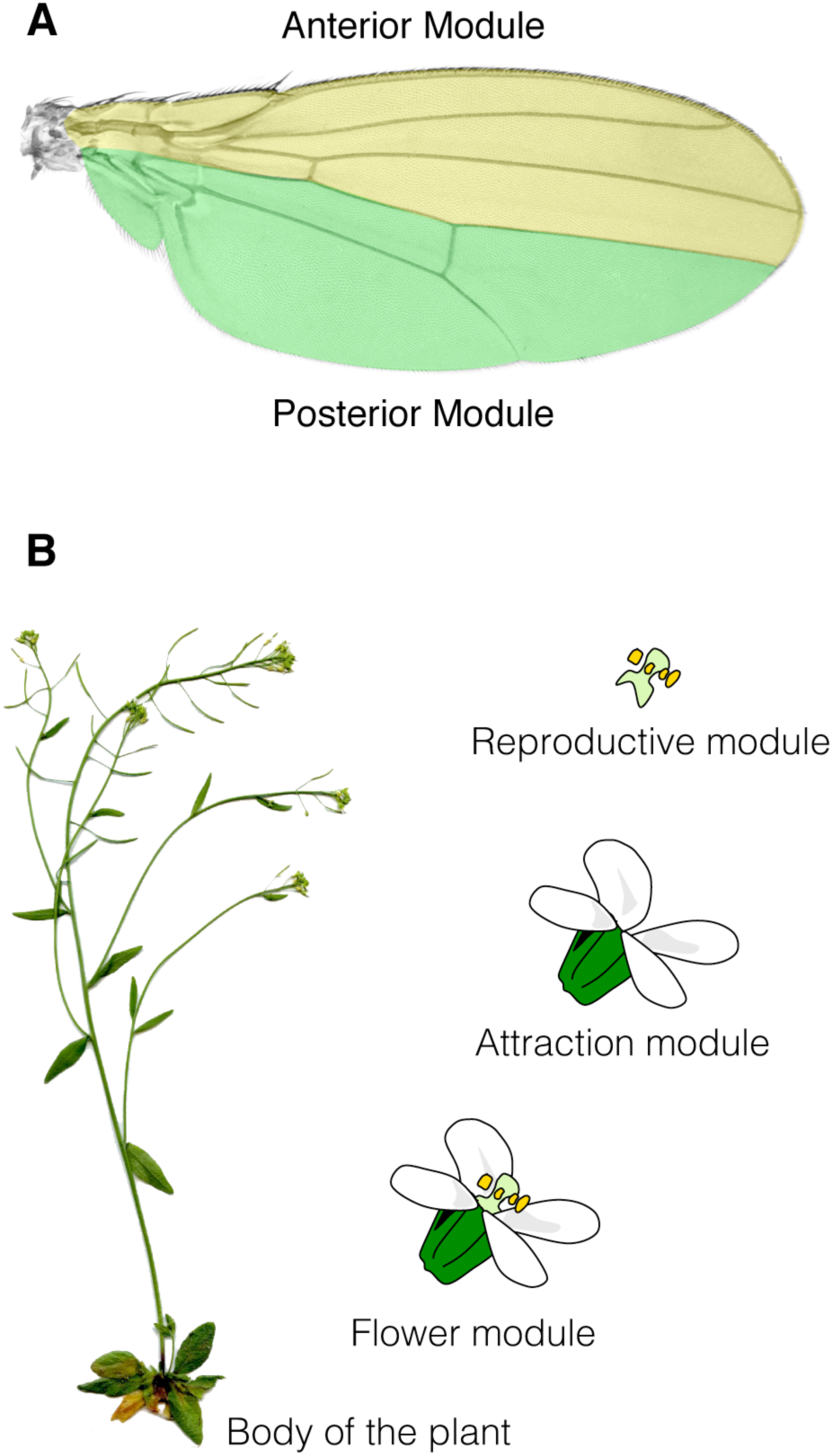
(A) Modules of the wing of insects illustrated in *Drosophila suzukii* (Diptera: Insecta). The wing comprises an anterior and a posterior module that derive from different developmental compartments. (B) Modules of flowering plants illustrated in *Arabidopsis thaliana* (Brassicales: Eudicots). Flowering plants are often divided into vegetative parts (roots, stems and leaves) and reproductive parts (flowers). In turn, the flower is divided into parts related to the attraction of pollinators and parts related to the pollination itself. Image credits: original photograph of the wing by Martin Hauser (Source: Wikimedia commons); original photograph of the plant by Lot Nature (Source: www.lotnature.fr); original drawing of the flower by Yvon Jaillais (Source: www.ens-lyon.fr/RDP/SiCE/Resources.html).

#### The body of angiosperms

The gross morphology of flowering plants shows often a modular division into vegetative parts (roots, stems and leaves) and reproductive parts (flowers) (Fig. 6B). It is common to find that these parts are repeated several times in a single plant (i.e. they are metameres), and hence, each one can be considered a module in itself. This distinction between vegetative and reproductive modules has its origin in the work of Raissa Berg on phenotypic pleiades (Conner and Lande 2014). Current research on plant modularity focuses on uncovering more precisely the contribution of genetic and developmental constraints, and of natural selection (in particular, that related to pollination), in the organization of phenotypic integration (e.g. Murren 2002; Rosas-Guerrero et al. 2011). In general, most of our knowledge on plant modularity comes from the study of flowers; in the literature reviewed, results suggest a division of the flower into functional and developmental modules, as well as into pollen-transfer and pollinator-attraction modules. The modularity of the flower has been recently reviewed in detail by Diggle (2014).

## 4. CONCLUSION

A modular organization of the body, with parts that develop, function, and evolve in a quasi-independent fashion, is common in both animals and plants. Modularity directly impacts the evolution of living beings because the relative integration within and between modules alters the evolvability and plasticity of organismal form. For this reason, a better understanding of the modular organization of living beings is essential to understand how their morphological variation is structured during evolution and development. The reviewed evidence suggests that our knowledge about morphological modularity is biased toward the study of mammals (in particular, *Homo* and *Mus*), whereas plants and arthropods are underrepresented despite having a well-defined modular body plan. Although the preferred object of study is the head of vertebrates, muscles and other soft tissues (except the brain) are systematically ignored in many studies; thus, reinforcing a bias toward hard tissues that downplays of the impact of muscles on shaping the head’s modularity. In general, studies on morphological modularity test only one hypothesis based, simultaneously, in two or more criteria. All in all, three out of four studies report the presence morphological modules, whereas fewer studies revealed that integration among parts was stronger than modularity. This finding indicates that, even though modularity is ubiquitous, the degree of modularity/integration varies depending on the morphological system studied, which it is expected if one considers a hierarchical organization of the body. Interestingly, factors explaining modularity and whole-system integration are used differentially in animals and plants. In animals, developmental and functional factors are used alike to explain the presence and the absence of morphological modules. In plants, function and environment (external factors) are used more frequently to explain a modular phenotype, while development and genes (internal factors) are used more frequently to explain a whole-phenotype integration. The findings of this systematic review identified some biases that must be overcome in order to reveal a whole new picture of how morphological modularity develops and evolve in complex living beings.

## ACKNOWLEDGEMENTS

I thank Diego Rasskin-Gutman and Rui Diogo for their feedback and discussions on modularity. This work would not have been possible without all the authors that made their articles available through public repositories and webpages. Special thanks to Maria Marín and Carles Felip-Leôn for their help in gathering full texts via institutional subscriptions to journals, and to Anna Loy, Erin Maxwell, Günter Wagner, John Maisay, and Pamela Diggle for kindly sending copies of their articles. This project has received funding from the European Union’s Horizon 2020 research and innovation programme under the Marie Sklodowska-Curie grant agreement No 654155.

